# STAT3 Serine phosphorylation is required for TLR4 metabolic reprogramming and IL-1β expression

**DOI:** 10.1101/2020.06.02.130716

**Authors:** Jesse Balic, Hassan Albargy, Kevin Luu, Francis J Kirby, W. Samantha N. Jayasekara, Finbar Mansell, Daniel J Gamara, Dominic De Nardo, Nikola Baschuk, Cynthia Louis, Fiachra Humphries, Katherine Fitzgerald, Eicke Latz, Daniel J Gough, Ashley Mansell

**Affiliations:** Centre for Innate Immunity and Infectious Diseases, Hudson Institute of Medical Research, Clayton, Victoria, 3168, Australia; Department of Molecular and Translational Sciences, Monash University, Clayton, Victoria, 3168, Australia; Centre for Cancer Research, Hudson Institute of Medical Research, Clayton, Victoria, 3168, Australia; Department of Anatomy and Developmental Biology, Monash Biomedicine Discovery Institute, Monash University, Victoria, 3800, Australia; Inflammation Division, The Walter and Eliza Hall Institute of Medical Research, 1G Royal Parade, Parkville, VIC 3052, Australia; Medical Biology, University of Melbourne, Parkville, Australia; Division of Infectious Diseases and Immunology, University of Massachusetts Medical School, Worcester, MA, USA; Institute of Innate Immunity, University Hospital Bonn, University of Bonn, Bonn, Germany; German Center for Neurodegenerative Diseases (DZNE), Bonn, Germany

## Abstract

Detection of microbial components such as lipopolysaccharide (LPS) by Toll-like receptor (TLR)-4 expressed on macrophages induces a robust pro-inflammatory response which has recently been shown to be dependent on metabolic reprogramming ^1, 2, 3, 4^. These innate metabolic changes have been compared to the Warburg effect (also known as aerobic glycolysis) described in tumour cells ^5, 6^. However, the mechanisms by which TLR4 activation leads to mitochondrial and glycolytic reprogramming remain unknown. Here we show that TLR4 activation induces a signalling cascade recruiting TRAF6 and TBK-1, while TBK-1 phosphorylates STAT3 on S727. Using a genetically engineered mouse model incapable of undergoing STAT3 Ser727 phosphorylation, we show both *ex vivo* and *in vivo* that STAT3 Ser727 phosphorylation is critical for LPS-induced glycolytic reprogramming, the production of the central immune-metabolite succinate and inflammatory cytokine production in a model of LPS-induced inflammation. Our study identifies non-canonical STAT3 activation as the crucial signalling intermediary for TLR4-induced glycolysis, macrophage metabolic reprogramming and inflammation.

## INTRODUCTION

TLR4 activation induces dramatic metabolic changes in macrophages via mitochondrial reprogramming, which is required to meet the rapid increase in demand for biosynthetic precursors for lipids, proteins, nucleic acids and the increased energy demand of the inflammatory state ^1, 2, 3, 4^. LPS-induced transcription of inflammatory cytokines and chemokines is also dependent on metabolic reprogramming ^2, 7^

Activated macrophages become more glycolytic, increase reactive oxygen species (ROS) production, and accumulate the tricarboxylic acid (TCA) cycle metabolite succinate redirecting it away from oxidative phosphorylation via the electron transfer chain (ETC). Recent studies have identified that the TLR downstream kinases TBK-1 and IKKε play a role in inducing glycolysis in immune cells ^8, 9^ via phosphorylation of Akt ^8^, although how these signals converge on, and orchestrate mitochondrial metabolic function remain unclear.

Metabolic signalling in response to elevated succinate concentration leads to the stabilization of hypoxia inducible factor-1α (HIF-1α), which positively regulates the prototypic inflammatory cytokine IL-1β and other HIF-1α-dependent genes involved in glycolysis ^10^. However, the molecular mechanisms of how membrane-bound TLRs communicate this signal to the mitochondria to alter mitochondrial function are also unclear.

STAT3 is a critical signalling molecule activated by immune cytokines resulting in phosphorylation on Tyr705 and activation of its activity as a transcription factor ^11^. In addition to this essential activity in the nucleus we, and others, have shown that a pool of Ser727 phosphorylated STAT3 translocates into the mitochondria where it affects mitochondrial metabolism and ROS generation ^12, 13^. In this study we interrogate the activation of STAT3 by TLRs as a mechanism to directly alter mitochondrial metabolism.

## RESULTS AND DISCUSSION

Activation of TLRs, with the exception of TLR3, sequester MyD88 to the receptor complex, initiating recruitment of the serine kinases IRAK1 and IRAK4 into the Myddosome complex ^9, 14, 15, 16, 17^. This complex subsequently interacts with TRAF6 to activate the canonical signaling pathway, resulting in the nuclear translocation of NF-κB and induction of the classic pro-inflammatory response ^18^. Alternatively, TLR3 and TLR4 also engage the adaptor TRIF leading to the activation of the serine kinase TBK-1 and subsequent phosphorylation of IRF3 leading to nuclear translocation and induction of IFNβ expression.

TRAF6 represents a major point of bifurcation of TLR signalling between canonical and non-canonical induction of inflammation. Structural analysis of TRAF6-binding partners show a conserved binding motif consisting of Pro-X-Glu-X-X-(aromatic or acidic residue) ^19^, required for interaction with downstream signalling proteins including Mal/TIRAP, TRIF, TRAM and STAT1^20, 21, 22, 23^. Given the homology between STAT proteins and the role for STAT3 in mitochondrial reprogramming, we identified highly conserved putative TRAF6-binding motifs within STAT3 (Fig.1a) and confirmed that the interaction between endogenous TRAF6 and STAT3 occurs within 10 minutes of LPS-stimulation in macrophages (Fig. 1b). To determine the importance of the putative TRAF6 binding sites in STAT3 we generated a series of mutant STAT3 constructs which were transiently expressed in 293T cells and bound to recombinant GST-tagged TRAF6 *ex vivo*. This data revealed selectivity for Glu100, i.e., STAT3 E100A completely abolished the interaction with TRAF6 which was not observed for the other putative interaction motifs (Fig 1c).

**Fig 1.**
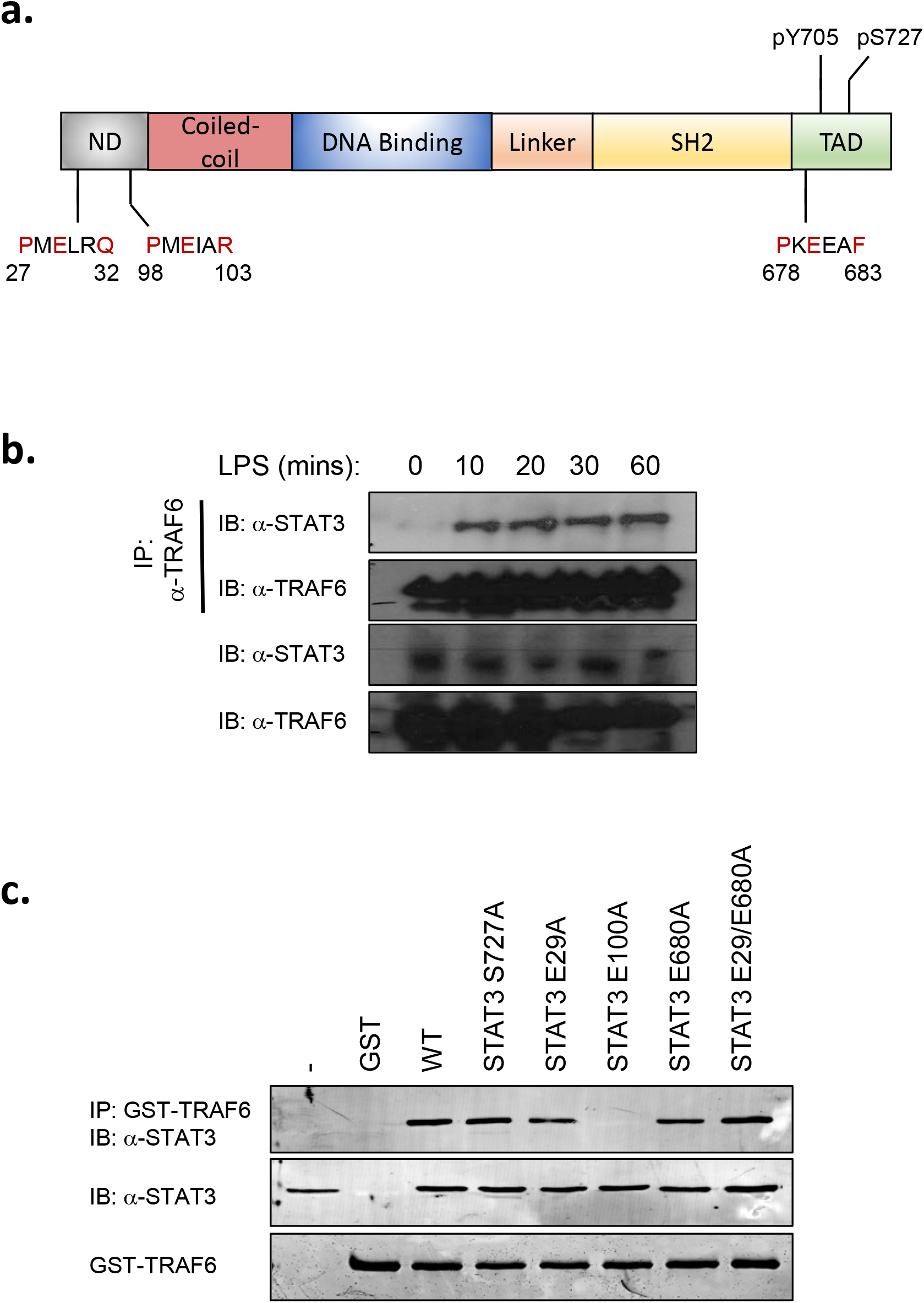
STAT3 directly interacts with TRAF6 following LPS stimulation. (**a**) Schematic representation of the putative TRAF6-binding domains and phosphorylation sites of STAT3. ND: N-terminal domain: TAD: transactivation domain. (**b**) BMDMs (1×10^7^ cells) were seeded in 10cm culture dishes and grown for 24 h prior to stimulation with LPS (1μg/ml) for indicated times. TRAF6 was immunoprecipitated from cellular lysates and probed with anti-TRAF6, or STAT3 antibody (n=3) (**c**) HEK293T cells (1×10^6^) were transfected with indicated plasmid vectors for 24 h. Cells were lysed and probed with recombinant GST-TRAF6 to immunoprecipitated interacting proteins, separated by SDS-PAGE and probed with anti-FLAG antibody (n=3).

Initially, we treated immortalized bone marrow-derived macrophages (iBMDMs) with IFNα as a well-known activator of STAT3. While we observed background Ser727 phosphorylated STAT3 in untreated cells that was endemic to iBMDMs, these cells did exhibit rapid increases in phosphorylation on both Ser727 and Tyr705 (Fig 2a) as expected. In contrast TLR ligands are reported to initiate phosphorylation of STAT3 on Tyr705 only after prolonged stimulation more consistent with secondary and indirect activation. To determine whether TLR activation induced rapid STAT3 phosphorylation, we challenged primary bone marrow-derived macrophages (BMDMs) with LPS and observed S727 phosphorylation within 20 mins of challenge, whereas Tyr705 phosphorylation was not detected even after 120 mins (Fig. 2b). To determine whether STAT3 Ser727 phosphorylation was induced by all TLRs we stimulated macrophages with Pam3Cys (TLR2), poly I:C (TLR3), Loxoribine (TLR7), CpG-DNA (TLR9), or IFNα as a positive control. All TLR agonists induced Ser727, but not Tyr705 phosphorylation of STAT3, whilst IFNα induced both pSer727 and pTyr705 as expected (Fig. 1c).

**Fig. 2.**
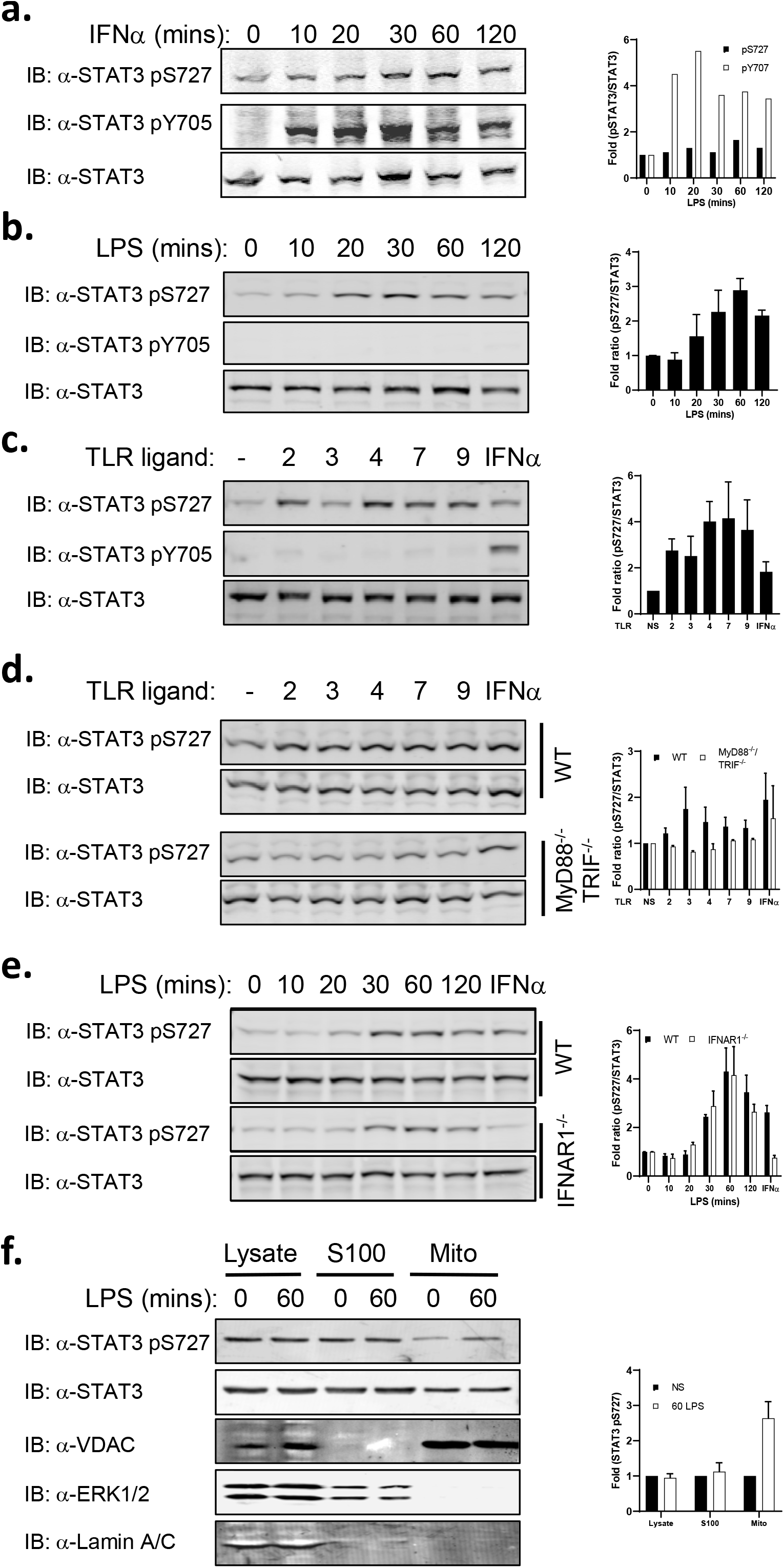
STAT3 undergoes non-canonical activation and mitochondrial localization following TLR stimulation. (**a**) Immortalized BMDMs (iBMDMs) (1×10^6^) were treated with IFNα (1000U) for indicated times, (**b**) BMDMs (1×10^6^ cells/well) were treated with LPS for indicated times, and (c) BMDMs were also treated with TLR agonists, Pam3Cys (TLR2, 100ng/ml), Poly I:C (TLR3, 500ng/ml), LPS (TLR4, 100ng/ml), Loxoribine (TLR7, 10mM), CpG-DNA (TLR9, 2.5μM) and IFNα (IFNAR1, 1000U) for 60 mins. STAT3 phosphorylation status was determined by immunoblot of cellular lysates with anti-STAT3, anti-STAT3 pS727 and anti-STAT3 pY705 antibodies as indicated (n=3). (**d**) MyD88^-/-^/TRIF^-/-^ iBMDM or (**e**) IRNAR1^-/-^ BMDMs were treated with TLR ligands (n=2) as described and immunoblot analysis with anti-STAT3 pS727 and anti-STAT3 antibodies of cellular lysates show ablated TLR-induced STAT3 Ser727 phosphorylation compared to IFNAR1-deficient BMDMs (Fig 3e), but not IFNα (n=2). (F) iBMDMs were treated with PBS or LPS (100ng/mL) for 1 hour and whole cell lysate, S100 or clean mitochondrial fractions isolated and separated by SDS-PAGE. Abundance of STAT3 and pS727 STAT3 in each fraction was determined as was fraction purity by probing with antibodies against Erk2 (S100), VDAC (mitochondria) or Lamin A/C (nucleus).

Stimulation of the TLR signalling pathway has at least two phases of response. The early response which is dependent upon the Myddosome complex components MyD88 and TRIF, leading to the activation of NF-κB and Interferon regulatory factor (IRF)-3 which drive the transcription of a suite of inflammatory genes including type I IFN ^18^; and the secondary response to secreted type I IFN. To confirm that STAT3 Ser727 phosphorylation is due to TLR activation directly and not as a consequence of autocrine cytokine or type I IFN signalling, we determined the phosphorylation of STAT3 on S727 in cells lacking the critical Myddosome factors MyD88 and TRIF or lacking the type I IFN receptor (IFNAR1). We show that LPS does not induce STAT3 phosphorylation on S727 in MyD88^-/-^/TRIF^-/-^ iBMDMs (Fig. 1d) but in unaffected in IFNAR1^-/-^-deficient macrophages (Fig 1e). Taken together these results demonstrate, for the first time, that STAT3 is directly recruited into the TLR signalling pathway via interaction with TRAF6, resulting in STAT3 Ser727, but not Tyr705 phosphorylation.

TLR-induced Ser727 phosphorylation of STAT3, in the absence of Tyr705 phosphorylation is consistent with the activation of the novel mitochondrial STAT3 activity that we have previously described in RAS transformed cancer cells ^12^. While the majority of STAT3 is present in the cytosol, a pool of STAT3 translocates into the mitochondria, causing metabolic reprogramming, and altering ROS production, dependent on pSer727 but independent of Tyr705 phosphorylation ^12^. We therefore, performed biochemical fractionation of LPS-treated macrophages (Fig. 1f) and demonstrate an enrichment of STAT3 pSer727 in mitochondrial fractions after 60 mins LPS stimulation. Taken together, these results demonstrate that STAT3 undergoes rapid TLR-induced Ser727, but not Tyr705 phosphorylation and an accumulation in the mitochondrial fraction of macrophages.

We next wished to identify the potential kinase responsible for TLR-induced STAT3 Ser727 phosphorylation. We have previously shown that the mitochondrial activity of STAT3 results in increased mitochondrial ROS (mtROS) production ^24^. Therefore, to identify potential kinases responsible for LPS-induced STAT3 Ser727 phosphorylation we used mtROS concentration as a functional readout of activity and screened a library of 355 kinase inhibitors (Supplementary Table 1).

Macrophages were pre-treated with inhibitors for 30 minutes prior to LPS challenge and mtROS production monitored every hour for 4 hours (Supplementary Fig. 1) using the mitochondrial superoxide probe MitoSOX. As a positive control we included the ROS scavenger N-Acetyl-L-cysteine (NAC) in all screens. This screening approach identified several classes of inhibitors capable of reducing LPS-induced mtROS production (Fig. 2a); including PI3K, mTOR and Inhibitor of IκB kinase (IKK) inhibitors. Interestingly, we also identified an inhibitor of tank binding kinase-1 (TBK-1) as a suppressor of LPS-induced mtROS production.

Importantly,IRAK1 and IRAK4 are serine kinases that interact with TRAF6 ^19^ and the MyDDosome whose kinase activity is critical for TLR-signalling and NF-κB activation ^14^. We found however, that a specific inhibitor of IRAK-1/4 activity had no effect on LPS-induced mtROS production, and therefore may not be responsible for STAT3 phosphorylation, acting as a specificity control for this screen.

This is consistent with the recent publication from Tan and Kagan ^9^ who showed that TRAF6 depleted macrophages are defective for TBK-1 recruitment to the Myddosome and induction of TLR glycolysis. Analysis of the TBK-1 sequence revealed a putative TRAF6 binding motif (human amino acids 223-228) suggesting the potential interaction between these proteins. Indeed, immunoprecipitation of endogenous protein show that TBK-1 directly interacts with TRAF6 following LPS stimulation in a time-dependent and transient manner (Fig 2b). This TBK-1-TRAF6 interaction was observed within 10 mins of LPS stimulation and was no longer detectable after 120 minutes post-stimulation. Our previous data demonstrate interaction between TRAF6 and STAT3 and led us to propose the formation of a TBK-1, TRAF6, STAT3 complex to enable TBK-1 phosphorylation of STAT3 in response to LPS challenge. Consistent with this we show that STAT3 interacted with phosphorylated TBK-1, with kinetics that are slightly offset from that of the TRAF6-TBK-1 interaction (Fig. 2c). Importantly, the immunoprecipitated STAT3 was phosphorylated on Ser727.

Whilst interaction studies and our mtROS inhibitor screen suggest that TBK-1 is the kinase upstream of STAT3 S727 phosphorylation in response to LPS stimulation we wanted to formally test this. We therefore examined STAT3 S727 phosphorylation in TBK-1-deficient macrophages. As can be seen in Figure 3d, we observed a consistent reduction in STAT3 S727 phosphorylation of approximately 50% when compared to WT cells, suggesting that TBK-1 is involved in STAT3 Ser727 phosphorylation. Previous studies have shown that TBK-1 and its closely related kinase IKKε are involved in TLR-induced glycolytic reprogramming ^8^ and that deletion of TBK-1 in IKKε^-/-^ macrophages reduced LPS-induced glycolysis that was dependent upon TRAF6 interaction suggesting that IKKε may play a role also in STAT3 phosphorylation.

**Fig. 3.**
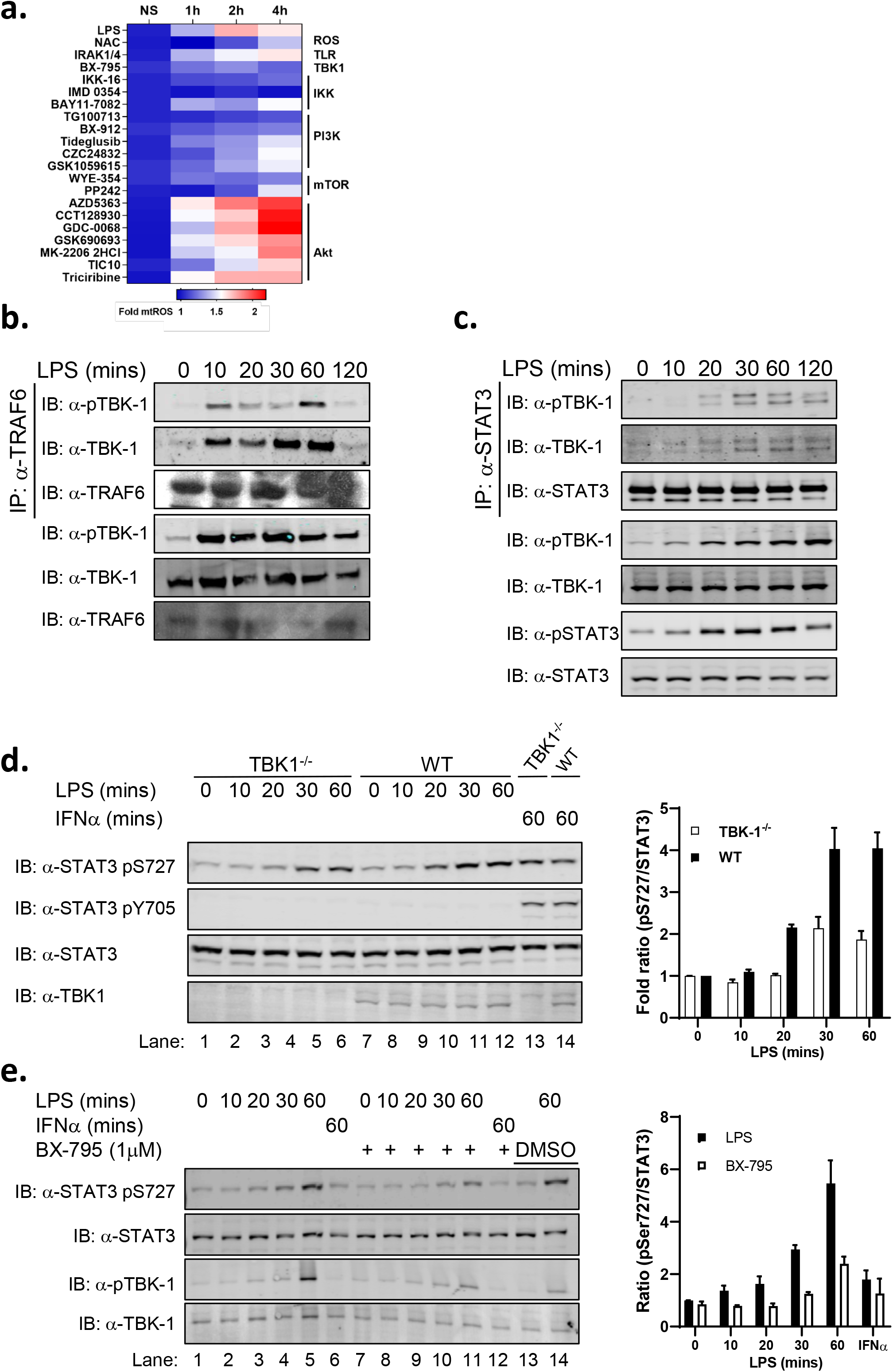
TBK-1 interacts with and plays a role in STAT3 Ser727 phosphorylation. (**a**) iBMDMs were seeded at 4×10^4^ cells per well in triplicate 24 h prior to pretreatment with kinase inhibitors (Y-axis; 500nM) for 30 mins. MitoSOX was added for 10 mins prior to challenge with LPS (100ng/ml; 0, 1, 2 and 4 h). LPS-induced production of superoxide by mitochondria was analysed by measuring oxidized MitoSOX fluorescence at 580nm. See also Supplementary Fig. **1a**. Cell lysates from LPS stimulated BMDMs (0-120 mins) were immunoprecipitated with (**b**) anti-TRAF6 and (**c**) anti-STAT3 antibodies. TBK-1 interaction with TRAF6 and STAT3 was identified by immunoblot with anti-TBK-1 and anti-TBK-1 pS172 antibodies respectively. (**d**) TBK-1-deficient and WT macrophages were treated with LPS or IFNa for indicated times and STAT3 phosphorylation determined in cellular lysates by immunoblot with indicated antibodies. TBK-1 expression was determined by immunoblot with anti-TBK-1 antibody. (**e**) BMDMs were pretreated with the TBK-1/IKKε (1μM) inhibitor BX-795 for 60 mins prior to LPS stimulation for indicated times. STAT3 and TBK-1 phosphorylation was determined by immunoblot with indicated antibodies. DMSO was added to lanes 13-14 as vehicle control. Immunoblot results are representative of at least three independent experiments.

We therefore pre-treated macrophages the inhibitor BX-795, at 1μM, at which concentration it is known to target both TBK-1 and IKKε ^25^, for 30 minutes prior to LPS stimulation. This pre-treatment completely reduced STAT3 Ser727 phosphorylation to background levels (Fig. 2e). These data suggest that the closely related kinases TBK-1 and IKKε are involved in STAT3 Ser727 phosphorylation. These findings are consistent with previous studies identifying interaction between overexpressed TRAF6 and TBK-1, and that these kinases may also play a role in downstream TLR signalling independent of their established role in mediating TLR-induced IRF3 phosphorylation and induction of IFNβ ^26^. In contrast we were unable to observe any direct interaction between STAT3 or TRAF6 with IKKε (data not shown. While PI3 kinase and Akt have previously been implicated in TLR-induced glycolytic reprogramming via TBK-1/IKKε ^8^, none of the seven Akt inhibitors screened in our study reduced LPS-induced mtROS (Fig 3a). We therefore examined whether the PI3 kinase inhibitor TG100713 could also inhibit LPS-induced STAT3 Ser727 phosphorylation and noted that this inhibitor had no effect upon LPS-induced phosphorylation (Supplementary Fig. 2). Therefore, while Akt has previously been identified as a downstream target of TBK-1/IKKε in TLR-induced glycolysis in dendritic cells ^8^, neither Akt nor PI3 kinase were implicated in TBK-1-mediated glycolysis in macrophages ^9^ and do not play a role in STAT3 S727 phosphorylation in macrophages. Together however, these studies identify a TRAF6, TBK-1/STAT3 signalling nexus leading to STAT3 S727 phosphorylation and mitochondrial localisation, providing a potential mechanism for the previously described role of TBK-1/IKKε signalling in TLR-induced glycolysis ^8, 9^.

Indeed, while non-canonical TBK-1 recruitment and phosphorylation was identified as critical in macrophage LPS-induced glycolysis ^9^, the role of Akt in macrophage signalling was not established akin to dendritic cells ^8^. Moreover, the role of these kinases may reflect cell specific signalling differences between macrophages and dendritic cells or reflect the kinetics of early and late glycolytic metabolic reprogramming pathways.

While the induction of glycolysis and metabolic reprogramming is increasingly recognized for its importance in the inflammatory state in macrophages, the mechanism of how TLRs promote this response is unknown. Our discovery identifies a molecular function for non-canonical kinase activity in TLR signalling via TBK-1 phosphorylation of STAT3, which subsequently translocates to the mitochondria.

Given that TBK-1 has been previously demonstrated to promote TLR-dependent glycolysis and that we have reported STAT3 Ser727-dependent metabolic changes, we hypothesized that the loss of STAT3 Ser727 phosphorylation would impede metabolic reprogramming. To address this, we performed a panel of metabolic analysis on primary peritoneal macrophages obtained from mice in which a serine to alanine (S727A) mutation was knocked into the endogenous STAT3 locus ^27^ abolishing Ser727 phosphorylation (STAT3 S727A; herein termed STAT3 SA). We used peritoneal exudate cells (PECs) for these studies because of the known requirement for STAT3 in growth factor signalling and differentiation (e.g. CSF-1, GM-CSF pathways) ^28^, moreover, PECs represent the local immunological environment, consisting predominately of myeloid cells ^29^. We found that STAT3 SA PECs have a significant defect in both resting glycolysis and the early glycolytic burst observed in wild-type macrophages following LPS-challenge as determined by assaying the extracellular acidification rate (ECAR) (Fig. 4a). Together with the previous observations in TBK-1/IKKε-deficient cells ^9^, these data support the concept that phosphorylation of STAT3 induces metabolic reprogramming.

**Fig. 4.**
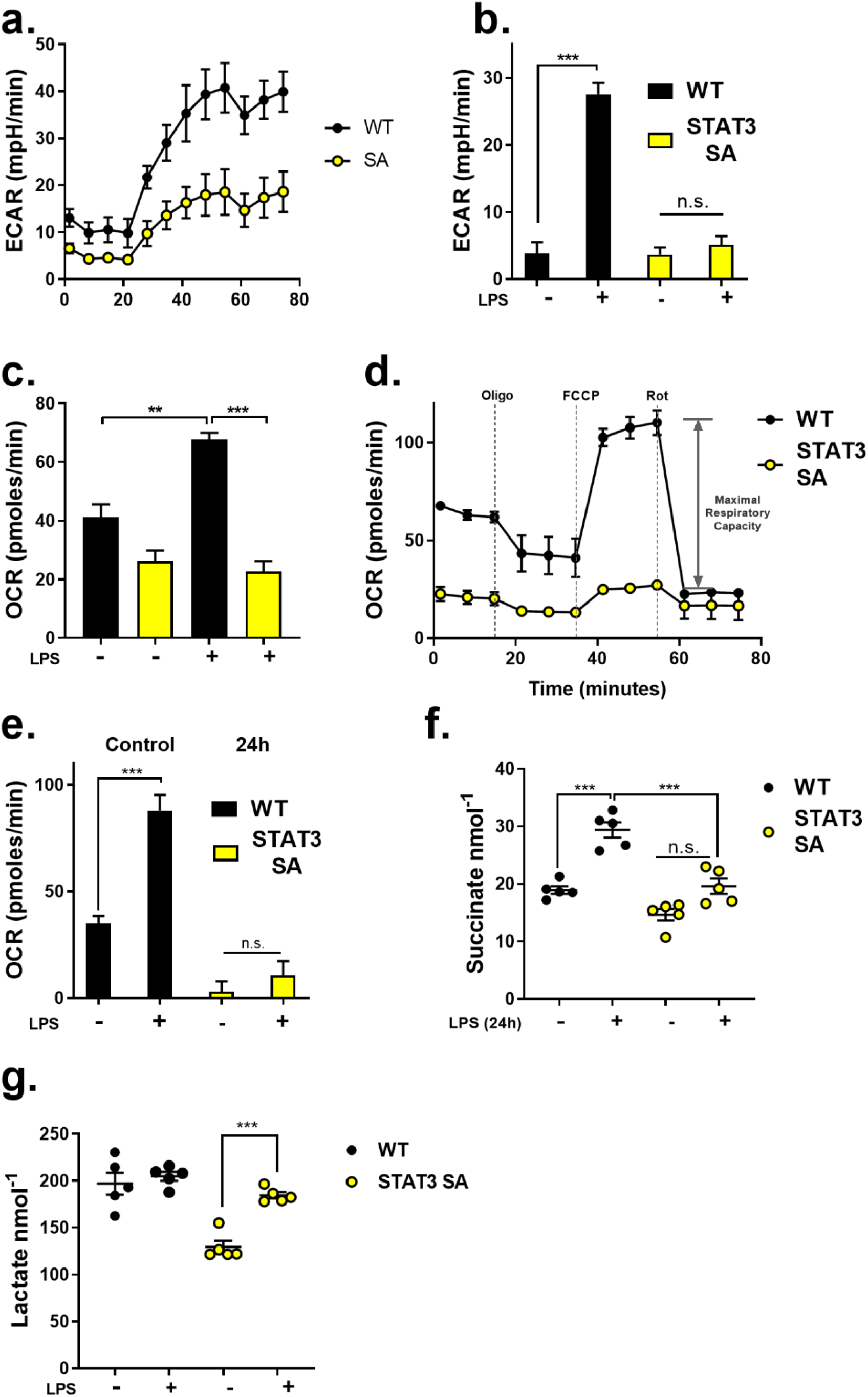
STAT3 Ser727 phosphorylation is critically for LPS-induced metabolic reprogramming and inflammation. Peritoneal macrophages (1×10^5^/well) obtained from WT vs STAT3 SA mice (n=3). (**a**) Real-time changes in the extracellular acidification rate (ECAR) of WT and STAT3 SA macrophages treated with LPS (**b-c**) WT and STAT3 SA peritoneal macrophages were stimulated with LPS or vehicle as indicated for 24 h. (**a-b**) The ECAR and oxygen consumption rate (OCR) were analyzed as indicators of oxidative phosphorylation and glycolysis, using a Seahorse XFp analyser. (**d**) Peritoneal macrophages were assayed for real-time changes in the OCR by sequential treatment with sequential treatment of cells with electron train chain inhibitors (oligomycin: adenosine triphosphate (ATP) synthase inhibitor; FCCP: H+ ionophore cyanide p-triflurmethooxyphenyl-hydrozone; and Complex I inhibitor rotenone). (**e**) Comparison of maximal respiratory capacity (MRC) between WT and STAT3 SA peritoneal macrophages stimulated with LPS or vehicle for 24 h. Results represent pooled macrophages obtained from 3 individual mice per genotype conducted in experimental triplicate, representing 3 independent experiments. Data presented as mean ± SEM, **p <0.001, ***p >0.0001, One-way ANOVA. (**f-g**) Peritoneal macrophages were seeded at 1×10^5^ cells/well and stimulated with LPS (100ng/ml) for 24 h prior to analysis of metabolites succinate and lactate produced in WT and STAT3 SA cultured supernatants. Data presented as mean ± SEM of 5 mice per genotype, ***p <0.001, Two-way ANOVA.

To determine whether STAT3 Ser727 phosphorylation is necessary for mitochondrial reprogramming, we exposed WT and STAT3 SA peritoneal macrophages to LPS for 24 hours and observed ablation of the ECAR and mitochondrial oxygen consumption rate (OCR) in STAT3 SA compared to WT cells (Fig. 4b and 4c). We next examined the functional capacity of the electron transport chain by determining the changes in OCR following sequential treatment of cells with mitochondrial electron transport chain (ETC) inhibitors. These data suggest that S727 phosphorylation of STAT3 is critical to elicit LPS-induced mitochondrial reprogramming and suggests STAT3 has parallel roles in LPS-induced metabolic reprogramming to that observed in tumour cells.

Consistent with what we have previously observed in cancer cells ^12^, the loss of STAT3 S727 phosphorylation leads to a diminished basal respiration rate. Moreover, STAT3 SA macrophages had a significant reduction in their maximal respiratory capacity compared to WT cells (Fig. 4d-e). These data mean whilst STAT3 SA PECs are viable and actively respiring, they are operating at close to their maximal capacity even in the absence of LPS stimulation. It should also be noted that the increase in basal respiration following a 24 h LPS-treatment of peritoneal macrophages we observe is the opposite of the LPS-mediated suppression of OCR observed in BMDMs, but is consistent with other studies on PECs ^30, 31^. This potentially reflects the different polarisation of these macrophage populations, where PECs display higher expression of M1 markers when compared to BMDMs ^32^. Indeed, in addition to the intrinsic metabolic differences in macrophages from diverse microenvironments, it has been suggested that the process of culturing BMDMs for 7 days in M-CSF culturing may push them towards M2 differentiation as compared to peritoneal macrophages ^33^. Therefore, while both cell types are display an intrinsic capacity to induce a potent inflammatory response to challenge, their metabolic responses differ.

Macrophage activation by LPS is accompanied by a remodelling of the tricarboxylic acid (TCA) cycle resulting in accumulating concentrations of TCA metabolites including succinate, which plays a critical role in mitochondrial reprogramming and inflammatory cytokine production. We found that while WT peritoneal macrophages treated with LPS for 24 hours displayed significantly increased succinate as expected. However, STAT3 SA macrophages failed to increase succinate concentrations (Fig. 3f). Our data also show that STAT3 SA macrophages have lower basal ECAR, which is increased in response to LPS but not to the magnitude, observed in WT macrophages. This is in line with our previous observation that mitochondria from STAT3 SA cells are defective in ETC activity ^12, 34^. Thus, STAT3 SA mitochondria may be more reliant on aerobic glycolysis that is consistent with the significant increase in the lactate concentration in STAT3 SA macrophages in response to LPS stimulation (Fig. 3g). However, the lactate concentration in LPS-treated STAT3 SA macrophages only ever approaches the lactate concentration observed in unchallenged WT macrophages. In addition, we do not observe any increase in the lactate concentration in WT macrophages in response to LPS stimulation. Given that ECAR is typically associated with lactate production this result appears somewhat counter-intuitive. However, it is important to note that ECAR also measures the export of CO2, hydration of H2CO3 and the dissociation to HCO3 and H^+^ from the respiratory chain which also contributes to the ECAR reading ^35^. Together, these data show that LPS induced STAT3 Ser727 phosphorylation is required for TCA cycle (succinate concentration) and OXPHOS (OCR) augmentation.

Previous studies have emphasized that enhanced succinate production is a critical regulator of the pro-inflammatory response via ETC-mediated mtROS production, the expression of IL-1β ^2, 10, 36^. Consistent with these observations we show that STAT3 SA macrophages generate significantly less *IL-1β* mRNA expression (Fig. 5a). We therefore, examined the kinetics and expression of cytokines in STAT3 SA PECs following LPS stimulation. Consistent with our mRNA data, while IL-1β expression increased steadily between 4-24 h of LPS challenge, IL-1β protein expression was significantly suppressed in STAT3 SA cell lysates compared to WT cells (Fig 5b). Furthermore, TNFα expression was reduced in STAT3 SA compared to WT cells (Fig 5c), but did continue to increase parallel to WT expression, while IL-6 concentrations were only significantly different at 24 h post-LPS (Fig 5d). Interestingly, whilst we observed increased IL-10 expression in unstimulated STAT3 SA macrophages, they did not respond to LPS with the increase in IL-10 production observed in WT cells (Fig 5e). These results demonstrate that STAT3 Ser727 plays a crucial role in inflammatory cytokine expression following LPS challenge. Importantly, these findings are consistent with previous studies characterizing metabolic reprogramming as integral to IL-1β production ^2, 10, 36^, while the temporal increase in cytokines such as IL-6 may be due to reduced autocrine induction due to reduced IL-1β or TNFα production. Interestingly it also suggests that STAT3 Ser727 phosphorylation may play a role in steady state IL-10 expression.

**Fig 5.**
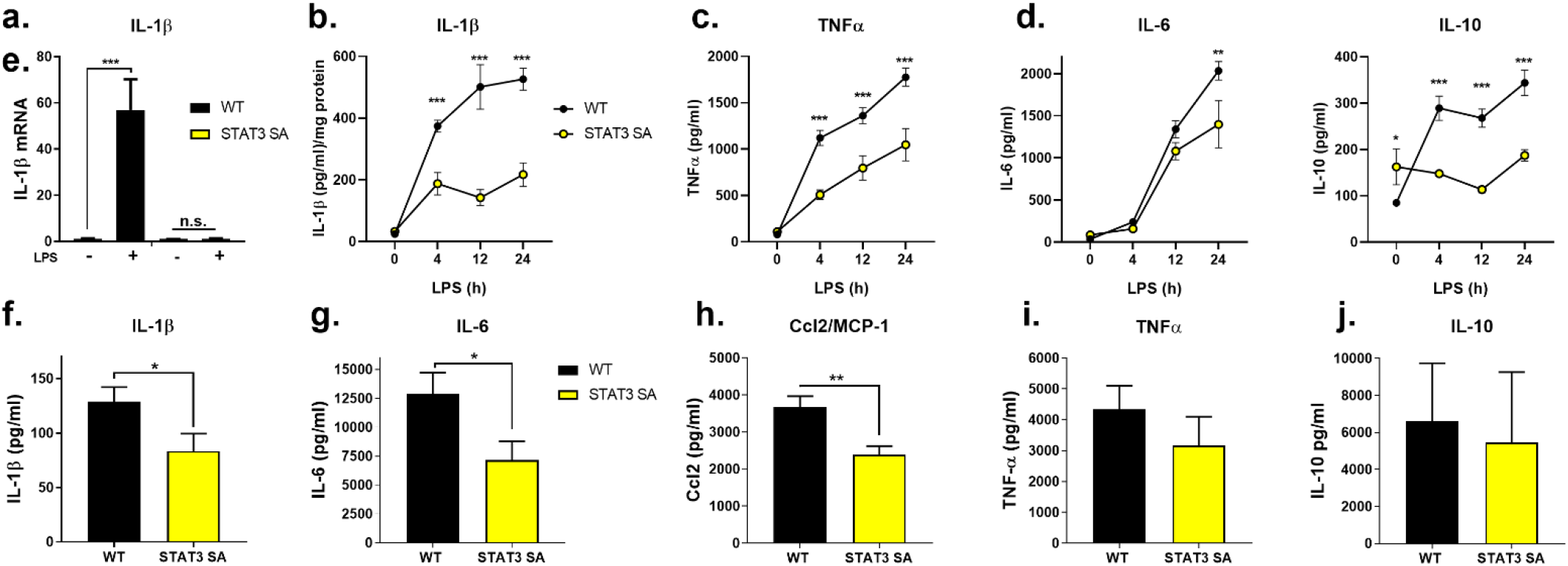
STAT3 S727 phosphorylation plays a role in LPS-induced cytokine expression. Peritoneal macrophages obtained from 3 mice per genotype were seeded at 5×10^5^ cells/well in and stimulated with LPS (100ng/ml) for 24 h prior to analysis IL-1β mRNA expression. Data presented as mean ± SEM of 5 mice per genotype, ***p <0.001, one-way ANOVA. Peritoneal macrophages generated from WT and STAT3 mice (3 mice per genotype) were seeded at 1.5×10^5^ in triplicate. Macrophages were stimulated for 4, 12 and 24 h with 100ng/ml of LPS and (**b**) cell lysates assayed for IL-1β and presented as IL-1β/mg of total protein, and cultured supernatants assayed for (**c**) TNFα, (**d**) IL-6, and (**e**) IL-10 by ELISA. Data presented are the pooled results of triplicate assays of three individual mice (n=9) per genotype, two-way ANOVA, *p < 0.05, **p < 0.01 and ***p < 0.001. WT and STAT3 SA mice were intraperitoneally treated with 10 mg/kg of LPS for 90 mins and serum analysed for (**f**) IL-1β, (**g**) IL-6, (**h**) MCP-1, (**i**) TNFα, and (**j**) IL-10 protein expression by ELISA or cytometric bead array (n=8/genotype, mean ± SEM, *p <0.05, **p <0.01, Students unpaired t test, two-tailed).

Increased aerobic glycolysis in macrophages plays a critical role in disease pathogenesis during endotoxemia ^10^. To investigate the role of STAT3 Ser727 phosphorylation in TLR-induced inflammation *in vivo*, we examined STAT3 SA mice in a model of LPS-induced sepsis. As can be seen in Figures 5f-i, STAT3 SA mice demonstrate significantly reduced IL-1β, IL-6 and Ccl2, but not TNFα, production in serum following acute LPS challenge, compared to WT mice. Consistent with our *in vitro* studies we observed no difference in IL-10 expression (Fig. 5j) which reflect the early response to LPS stimulation, whereas IL-10 expression may be delayed. Taken together, these results highlight the specificity of Ser727 phosphorylated STAT3 in mediating inflammatory gene induction.

Increasing evidence positions metabolic reprogramming as a key event in the inflammatory response following TLR activation. Importantly, this study identifies, for the first time, the molecular mechanism by which TLR signalling communicates with the mitochondria. We have demonstrated that TBK-1 contains a putative TRAF6 binding site allowing its direct recruitment to the TLR signalling pathway, facilitating TBK-1-mediated STAT3 Ser727 phosphorylation and translocation to the mitochondria. Macrophages unable to undergo STAT3 Ser727 phosphorylation display impaired TLR-induced glycolytic reprogramming, reduced pro-inflammatory metabolite production and diminished inflammation. This discovery parallels observations made in cancer cells, which further enhances the concept that inflammatory macrophages induce the Warburg effect to engender a pro-inflammatory phenotype. As such, STAT3 is not only a central immune regulatory transcription factor, but additionally rapidly orchestrates innate immune mitochondrial reprogramming and inflammatory cell metabolism via non-canonical signalling.

## METHODS

### Bone Marrow-Derived Macrophage (BMDM) Isolation and Cell Culture

Wild type, IFNAR1^-/-^ (kind gift from Prof Paul Hertzog, Hudson Institute), TBK-1^F/F^, TBK-1 ^F/F^/Vav-iCre BMDMs were differentiated in DMEM containing 30% M-CSF conditioned media obtained from supernatants of L929 fibroblasts (containing M-CSF), centrifuged to remove cell debris (5 min, 300g) and filtered through a 0.22μm filter. Leg bones of WT mice were surgically removed and cleaned bones were cut with scissors and flushed with sterile PBS via a syringe. Bone marrow suspension was passed through a 70mm cell strainer to remove clumps and cells cultured in low-adherence 10 cm tissue culture plates in L929 supplemented 10% FCS, DMEM with added L-glutamine at 37°C, 5% CO2 for 7 days. Cells were supplemented with a further 5ml of L929 conditioned FCS/DMEM on day 3. Cells were removed from tissue culture plates with gentle scrapping and seeded at desired densities in 1% FCS/DMEM, supplemented with L-glutamine 24 h prior to stimulation or treatment.

Immortalized Wild type and MyD88^-/-^/TRIF^-/-^ BMDMs were generated from indicated mice with J2 recombinant retrovirus carrying v-myc and v-raf oncogenes ^37, 38^.

All cell lines were confirmed by STR profiling prior to experimentation. HEK293T cells were obtained from ATCC were grown in10% FCS in DMEM supplemented with L-glutamine and grown in humidified 5% CO2 at 37°C.

### Immunoprecipitation of TRAF6 and STAT3

To determine the binding interface between STAT3 and TRAF6 we generated a panel of STAT3 mutants by site directed mutagenesis of E28, E100, E680 or E28 and E680 using the following primers E28A: F 5’-CAGTGACAGCTTCCCATGGCGCTGCGGCAGTTTC; R 5’-GAAACTGCCGCAGCGCCATGGGAAGCTGTCACTG, E100A: F 5’-CTTGAGAAGCCAATGGCGATTGCCCGGATTGTG; R 5’-CACAATCCGGGCAATCGCCATTGGCTTCTCAAG, E680A: F 5’-CACAATCCGGGCAATCGCCATTGGCTTCTCAAG, R 5’-CTTGAGAAGCCAATGGCGATTGCCCGGATTGTG. The resultant FLAG-tagged STAT3 constructs were transfected into 293T using lipofectamine 3000 (Thermo Fisher Scientific). 48 hours after transfection cells were lysed (50mM Tris, pH 7.4, 1.0% Triton X-100, 150mM NaCl, 1mM EDTA, 2mM Na3VO4, 10mM NaF, 1mM PMSF and protease cocktail inhibitor (Roche)) and centrifuged at 18,000g for 5 mins (4°C) to remove debris. GST tagged TRAF6 C-domain was cloned into pGEX-4T-3 and expressed in BL21(DE3) bacteria as described previously ^21^. Briefly, TRAF6 transformed BL21 bacteria were cultured in Luria Broth supplemented with ampicillin (100ug/ml) and when an OD600 of 0.6 reached TRAF6 expression was initiated by incubation with 500 μM Isopropyl β-d-1-thiogalactopyranoside (IPTG) for 16 hours at 16 °C. Bacteria was pelleted by centrifugation and lysed by incubation in 50 mM Tris, 150 mM NaCl at pH 7.6 with complete protease inhibitors (Roche) (10ml per g of bacteria) and sonicated at 4 °C for 30 second bursts with 30 seconds between for a total of 10 cycles. Bacterial lysate was clarified by centrifugation at 40,000 xg for 45 min at 4°C and the clarified protein bound to glutathione agarose beads (Thermo Fisher Scientific) at 4 °C for 4 hours with rotation. Beads were washed in 10x bead volume of lysis buffer three times. Protein induction and TRAF6-bound beads were assessed by SDS-PAGE and Coomassie blue staining of a protein of ~63kDa. To determine STAT3 binding to recombinant TRAF6, 20μl of GST-TRAF6 beads were added to equivalent, clarified lysate from transfected 293T cells described above. Binding was performed at 4 °C for 4 hours with rotation. Beads were washed in 10x bead volume of buffer 50mM Tris, pH 7.4, 1.0% Triton X-100, 150mM NaCl, 1mM EDTA, 2mM Na3VO4, 10mM NaF, 1mM PMSF and protease cocktail inhibitor (Roche)) three times for 5 minutes each with rotation. After the final wash supernatant was removed and beads resuspended in Laemmli buffer, boiled for 5 minutes and, together with input protein sample separated by SDS-PAGE. Gels were transferred to nitrocellulose, blocked in odyssey blocking buffer and incubated with indicated primary antibodies. Equivalent GST-TRAF6 loading was determined by Coomassie blue staining of SDS-PAGE gels following transfer.

### Immunoblotting of STAT3 phosphorylation

To determine the phosphorylation status of STAT3, BMDMs (1×10^6^) cells were seeded in 6-well plates 24 h prior to stimulation with agonists (IFNα, 1000U; Pam3Cys, 100ng/ml; poly I:C, 10μg/ml; LPS, 100ng/ml; Loxoribine, 500μM and CpG-DNA, 500nM) for indicated times. pBMDMs were lysed in a modified RIPA lysis buffer [50mM Tris (pH 7.4), 150mM NaCl, 1mM EDTA, 1% (v/v) Triton X-100, 0.3% (w/v) sodium deoxycholate, 0.3% (w/v) SDS supplemented with protease and phosphatase inhibitors (Roche). Protein concentration was determined using the *DC* Protein Assay (Bio-Rad). Samples were reduced and boiled in Laemmli buffer (Bio-Rad) containing 10% (v/v) beta-mercaptoethanol, resolved by SDS-PAGE, transferred onto PVDF (EMD Millipore), blocked in Odyssey Blocking Buffer (LI-COR) and incubated with the following primary antibodies overnight: pY705-STAT3, pS727-STAT3 and STAT3 (Cell Signaling Technology). Membranes were then probed with the appropriate IRDye conjugated secondary antibodies: anti-rabbit IgG (ThermoFisher Scientific), anti-mouse IgG (Rockland Immunochemicals), anti-rat IgG (Rockland Immunochemicals). Membranes were scanned using an Odyssey ^®^ Infrared Imaging System. To determine the role of kinases in STAT3 phosphorylation, BMDMs were pretreated with either 1μM BX-795 (Caymen Chemicals) or 1μM TG10073 (Selleck Chemicals) for 60 mins prior to LPS challenge and assayed as described above.

### Mitochondria isolation

Pure mitochondria were isolated from macrophages (iMAC) as previously published (Gough 2009) with minor amendments. 5×10^7^ – 1×10^8^ iMACs were collected by scraping with a rubber policeman, washed twice and resuspended in 5x the pellet volume of buffer A (20 mM HEPES pH 7.6, 220 mM mannitol, 70 mM sucrose, 1 mM EDTA, 0.5 mM phenylmethylsulfonyl fluoride (PMSF) and 2 mg/ml bovine serum albumin (BSA)]. Cells were incubated on ice for 15 min to facilitate cell swelling before being subjected to nitrogen cavitation under 200 PSI of pressure for 5 minutes (PARR instruments). Cell homogenate were centrifuged at 800 *g* for 10 min at 4°C and the mitochondria containing supernatant retained and centrifuged at 10,000 *g* for 20 min at 4°C. The supernatant representing the crude mitochondrial fraction was resuspended in 1 ml Solution B (20 mM HEPES pH 7.6, 220 mM mannitol, 70 mM sucrose, 1 mM EDTA, 0.5 mM PMSF) and loaded on top of a stepwise Percol gradient comprised of 1mL 80% Percol/balance solution A, 4.5mL 56% Percol/balance solution A and 4.5mL 23% Percol/balance solution A. Gradients were centrifuged at 65,000g for 45 minutes and mitochondria isolated from the junction of the 56 and 23% layers. Mitochondria were washed twice in solution B and mitochondrial protein content determined by adding 1 μl of mitochondrial suspension to 600 μl of 50 mM Tris pH 7.4, 0.1% (w/v) SDS and measuring the absorbance at 280nm and 310nm. Mitochondrial protein concentration in mg/ml is given by (A_280nm_-A_310nm_)/1.05 x 600 (PMID175068).

S100 fractions were prepared by collecting the supernatant after the initial 10,000g crude mitochondrial isolation centrifugation which was centrifuged at 100,000g in a Beckman benchtop ultracentrifuge. Protein concentration was determined by Bradford assay. Equivalent protein was resolved through SDS-PAGE and protein content and fractionation purity determined by western blot with antibodies against the indicated proteins.

To assess STAT3 mitochondrial enrichment, 2 x 10^7^ cells/ml into 24-well tissue culture plates in 900 μl of THP1-conditioned media. THP1 cells were then stimulated with Pam3Cys (100 ng/ml) or LPS (100 ng/ml) for indicated times. THP1 cells were pelleted at 300g for 3 mins then resuspended in 10 ml of chilled PBS. The cells were lysed in a 45 ml cell disruption vessel (Parr Instrument Company) using nitrogen cavitation at 350 psi for 1 minute. The resulting lysates were collected and using the Mitochondria Isolation Kit (Miltenyi Biotech) used to isolate mitochondria as per manufacturer’s instructions. Briefly, cells were pelleted at 2000 rpm for 10 mins and then resuspended in 1 ml of lysis buffer. 9 ml of 1x separation buffer was added to the lysis buffer and 50 μl of α-Translocase of the Outer Mitochondrial Membrane 22 (TOM22) microbeads were used to label mitochondria and incubated for 60 minutes. LS columns were placed into a MidiMACS separation unit and the columns were rinsed 3 times with separation buffer. Lysates were separated through the columns and washed three times with separation buffer. Additional buffer was added and flushed out by placing the plunger into the column. The mitochondrial suspension was further centrifuged at 13,000 rpm for 2 mins and the supernatant decanted with 100 μl of SDS-loading buffer added to the pellet and heated at 95°C for 5 minutes prior to analysis of proteins by immunoblot.

### STAT3 SA Mice

STAT3 S727A (SA/SA) mice were generated as previously described ^27^. C57BL/6J wild type (WT) and STAT3 SA mice were maintained at the Monash Medical Centre Animal Facility under specific pathogen-free conditions in accordance with Australian Government and Monash University. Female and male mice were used for both genotypes at 6-12 weeks of age.

### Peritoneal Macrophage Isolation and Cell Culture

Peritoneal cells were collected via peritoneal lavage with 5ml of cold sterile PBS supplemented with 5mM EDTA. Cells were centrifuged (300g, 5 mins) and allowed to adhere to tissue culture plates for 1 h in 10% FCS, DMEM and 2mM L-glutamine. Cells were washed three times with sterile PBS after 180 mins to remove non-adherent cells and grown overnight at 37°C, 5% CO2in 1% FCS/DMEM supplemented with 2mM L-glutamine prior to experimentation ^29^.

### Gene Expression Analysis by Quantitative Real-Time PCR and Cytokine Detection

Total RNA from WT and STAT3 SA peritoneal macrophages was isolated using the RNAeasy Isolation Kit (Qiagen) and reverse transcribed with random hexamers (Life Technologies) using Moloney murine leukemia virus reverse transcriptase (Promega) according to manufacturers’ instructions. mRNA was quantified with SYBR reagents (ABI) using primer pairs targeting *il1β* (Forward 5;-CAACCAACAAGTGATATTCTCCATG-3’, Reverse 5’-GATCCACACTCTCCAGCTGCA-3’), and 18S (Forward 5’-GTAACCCGTTGAACCCATT-3’, Reverse 5’-CGAATCGAATCGGTAGTAGCG-3’). Relative mRNA expression was analysed using the comparative CT method, normalizing genes of interest to the 18S housekeeping gene and fold gene induction calculated relative to expression in control samples.

Quantification of cytokines secreted from peritoneal macrophages and serum, and peritoneal macrophages cellular lysates were conducted using ELISA kits from R&D Systems or Cytokine Bead Array (BD Biosciences) according to manufacturers’ instructions.

### Monitoring Fluorescence Mitochondrial ROS production by LPS

Analysis of kinase inhibition of LPS-induced mitochondrial ROS production was modified from an earlier method by Wojtala et al ^29^. iBMDMs (4×10^4^/ml) were seeded in triplicate in black microtest Optilux 96-well plates (BD Falcon) in 1% FCS/DMEM and L-glutamine media for 4 h and then replaced with phenol-red free 1% FCS/DMEM and L-glutamine 20 h prior to 30 mins pretreatment or not with kinase inhibitors (500nM). Macrophages were incubated with a final concentration of 1μM MitoSox (Thermo Fischer) for 10 mins before treatment with LPS (100ng/ml). Plates were analysed for fluorescence emission as a marker of mtROS production (excitation/emission 510/580nm) every 60 mins for 4 h (ClarioStar Plate Reader, BMG) and expressed as the fold induction compared to nontreated control macrophages.

### Seahorse Assay and Metabolite Analysis

The extracellular acidification rate (ECAR) and oxygen consumption rate (OCR) of WT and STAT3 SA peritoneal macrophages were measured with a Seahorse XF Analyzer. To measure ECAR and OCR in real-time, cells were isolated from mice, seeded at 1×10^5^ cells/well in 8-well miniplate format and allowed to adhere overnight. Cells were washed twice in media and treated with LPS (100ng/ml) for 24 h. One hour prior to reading cells were washed twice with, and then cultured in Seahorse XF base medium supplemented with 1mM pyruvate, 2mM L-glutamine and 10mM glucose in an incubator without CO2. ECAR and OCR were measured under basal conditions prior to sequential treatment of cells with electron train chain inhibitors 1 μM oligomycin, 1.5μM FCCP-cyanide p-tribluromethoxyphenyl-hydrazone, and 1μM antimycin A and rotenone. In experiments examining real-time induction of ECAR by LPS, cells were isolated and seeded as described above, however following basal measurement of ECAR, cells were challenged with LPS (100ng/ml) and ECAR measured for 180 mins. Data represents mean ± SEM of triplicate wells from at least 3 independent mice.

Cultured supernatants from LPS-treated WT and STAT3 SA peritoneal macrophages were also assayed by Succinic Acid Colormetric Assay Kit (Biovision) and Lactate-Glo Assay (Promega) for succinate and lactate metabolite concentrations respectively according to manufacturers’ instructions.

### LPS-induced Model of Sepsis

LPS-induced sepsis model in mice was approved by the Monash Medical Centre Animal Ethics Committee. Sepsis was induced in male and female wild type (WT) and STAT3 SA mice (aged 6-14 weeks) following i.p. injection with10 mg/kg in a total volume of 100μl of LPS (E. coli 055:B5 ultrapure; Invivogen). Mice were culled after 90 mins and serum collected via cardiac puncture for measurement of serum cytokines.

## QUANTIFICATION AND STATISTICAL ANALYSIS

Statistical analyses were conducted using specific statistical tests as indicated in the figure legends using GraphPad 8.01 software for each experiment. Data are represented as the mean ± standard error of the mean as indicated in the figure legends which includes the biological and experimental replicates. Significance is depicted with asterisks on graph as follows: *p < 0.05, **p < 0.01 and ***p < 0.001.

## ACKNOWLEDGEMENTS

This work was supported by the Victorian State Government Operational Infrastructure Scheme. DJG is supported by a mid-career fellowship from the Victorian Cancer Agency (MCRF19033) and grants from the United States Department of Defence (CA150132) and the Cancer Council Victoria (GNT1145028). HA is supported by a scholarship from the College of Applied Medical Sciences, Shaqra University. The authors also wish to thank Dr Rebecca Smith with editorial assistance in preparing this manuscript.

## AUTHOR CONTRIBUTIONS

A.M, D.J.G. and EL conceived and designed the concept and experiments. J.B., H.A., K.L, F.J.K, W.S.N.J, F.M., D.J.G, N.B., and D.J.G designed, conducted and interpreted experiments. D.D.N, C.L., K.A.F, F.H and E.L. contributed reagents and the manuscript was written and edited by A.M. and D.J.G.

## DATA AVAILABILITY STATEMENT

The data that supports the findings of this study are available from the corresponding author upon reasonable request.

## FIGURE LEGENDS

**Supplementary Figure 1.**
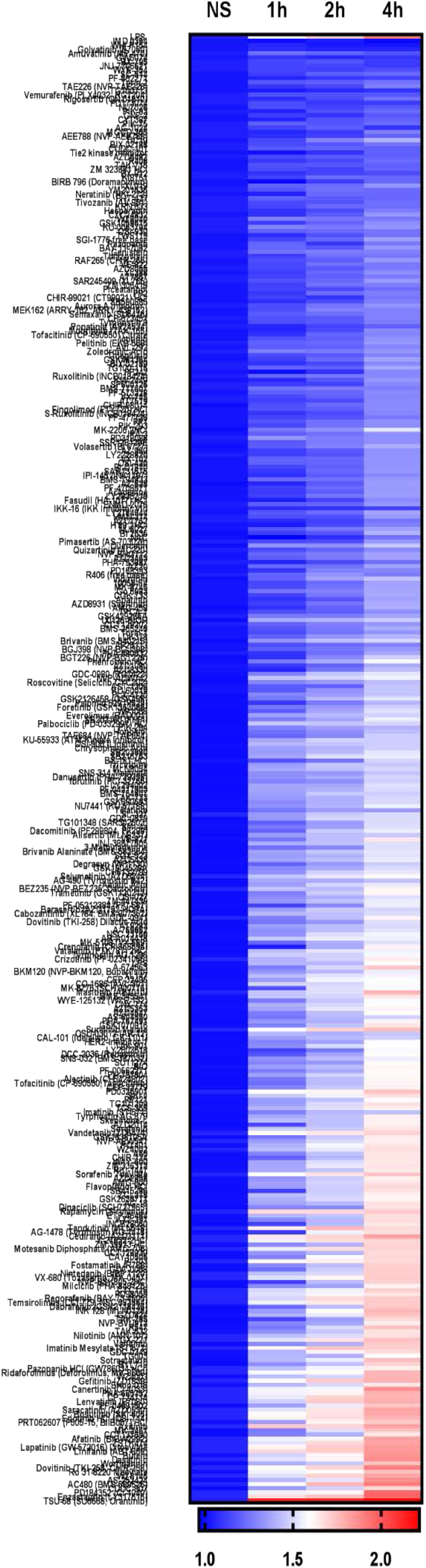

**Supplementary Figure 2.**
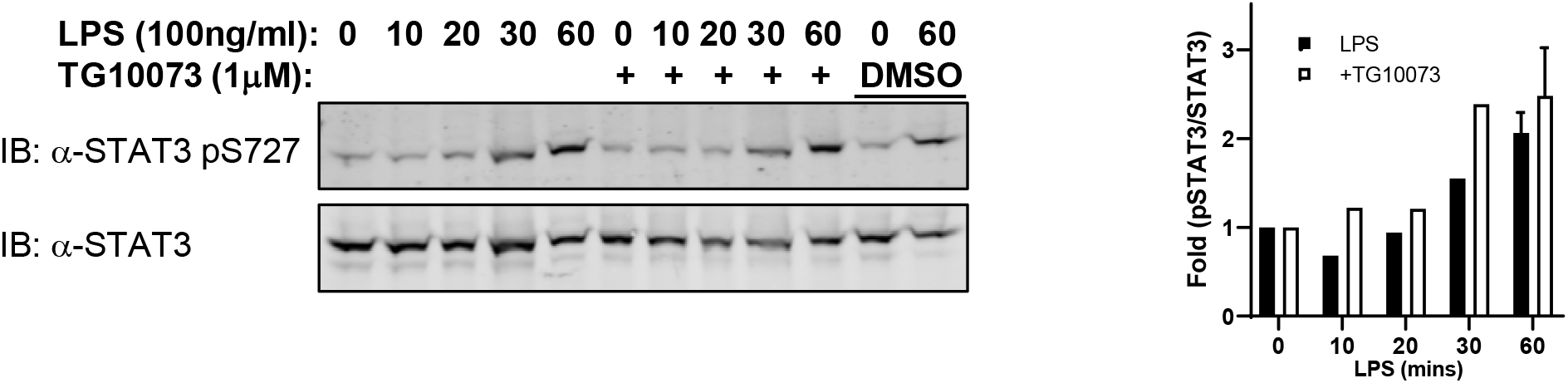

